# The DOE JGI Metagenome Workflow

**DOI:** 10.1101/2020.09.30.320929

**Authors:** Alicia Clum, Marcel Huntemann, Brian Bushnell, Brian Foster, Bryce Foster, Simon Roux, Patrick P. Hajek, Neha Varghese, Supratim Mukherjee, T.B.K. Reddy, Chris Daum, Yuko Yoshinaga, Rekha Seshadri, Nikos C Kyrpides, Emiley A. Eloe-Fadrosh, I-Min A. Chen, Alex Copeland, Natalia N. Ivanova

**Author notes:** First Author and Second Author contributed equally to this work. Author order was determined randomly. Address correspondence to Alicia Clum,.

## Abstract

The DOE JGI Metagenome Workflow performs metagenome data processing, including assembly, structural, functional, and taxonomic annotation, and binning of metagenomic datasets that are subsequently included into the Integrated Microbial Genomes and Microbiomes (IMG/M) comparative analysis system (I. Chen, K. Chu, K. Palaniappan, M. Pillay, A. Ratner, J. Huang, M. Huntemann, N. Varghese, J. White, R. Seshadri, et al, Nucleic Acids Rsearch, 2019) and provided for download via the Joint Genome Institute (JGI) Data Portal (https://genome.jgi.doe.gov/portal/). This workflow scales to run on thousands of metagenome samples per year, which can vary by the complexity of microbial communities and sequencing depth. Here we describe the different tools, databases, and parameters used at different steps of the workflow, to help with interpretation of metagenome data available in IMG and to enable researchers to apply this workflow to their own data. We use 20 publicly available sediment metagenomes to illustrate the computing requirements for the different steps and highlight the typical results of data processing. The workflow modules for read filtering and metagenome assembly are available as a Workflow Description Language (WDL) file (https://code.jgi.doe.gov/BFoster/jgi_meta_wdl.git). The workflow modules for annotation and binning are provided as a service to the user community at https://img.jgi.doe.gov/submit and require filling out the project and associated metadata descriptions in Genomes OnLine Database (GOLD) (S. Mukherjee, D. Stamatis, J. Bertsch, G. Ovchinnikova, H. Katta, A. Mojica, I Chen, and N. Kyrpides, and T. Reddy, Nucleic Acids Research, 2018).

**IMPORTANCE:** The DOE JGI Metagenome Workflow is designed for processing metagenomic datasets starting from Illumina fastq files. It performs data pre-processing, error correction, assembly, structural and functional annotation, and binning. The results of processing are provided in several standard formats, such as fasta and gff and can be used for subsequent integration into the Integrated Microbial Genome (IMG) system where they can be compared to a comprehensive set of publicly available metagenomes. As of 7/30/2020 7,155 JGI metagenomes have been processed by the JGI Metagenome Workflow.

## INTRODUCTION

Metagenomics, the study of the genetic content of natural microbial communities, provides a wealth of information about the structure and dynamics, perturbation, and resilience of ecosystems. Many tools are available for processing and analyzing metagenomic datasets including metaSPAdes(1) and MEGAHIT(2) for assembly, Prokka(3) and MG-RAST(4) for annotation and Kraken 2(5) for taxonomic identification, as well as integrated workflows such as SqueezeMeta(6) and MGnify(7). Here we present a metagenome workflow developed at the JGI which generates rich data in standard formats and has been optimized for downstream analyses ranging from assessment of functional and taxonomic composition of microbial communities to genome-resolved metagenomics and identification and characterization of novel taxa. This workflow is currently being used to analyze thousands of metagenomic datasets in a consistent and standardized manner.

## RESULTS

The DOE JGI Metagenome Workflow aims to provide consistently processed metagenome data in standard formats suitable for a wide variety of analyses and interpretations across many studies and environmental samples. The workflow performs multiple quality checks and artifact removal, and provides a variety of summary statistics to assist users with the assessment of data quality and consistency. We illustrate the workflow using microbiomes from the Loxahatchee Nature Preserve in the Florida Everglades(8) as an example. In this follow-up study, sediment samples were collected and DNA was isolated by the students of Boca Raton Community High School, Boca Raton, from 4 different sites in the Loxahatchee Nature Preserve with 5 replicates at each site as previously described. DNA isolated from these samples was sequenced at the JGI using Illumina NovaSeq and standard library and sequencing protocols (Kapa HyperPrep library preparation kit, see Methods). Raw 2×150 reads were then processed by the DOE JGI Metagenome Workflow. The metadata for these samples can be found in Genomes OnLine Database (GOLD) (9) using GOLD study Gs0136122. Raw reads, as well as intermediate results and final assembly and annotation data can be found in the JGI Data Portal (https://genome.jgi.doe.gov) usingJGI sequencing project identifiers linked to the GOLD study and Integrated Microbial Genome (IMG)(10) taxon identifiers provided in Table 1.

**TABLE 1.**
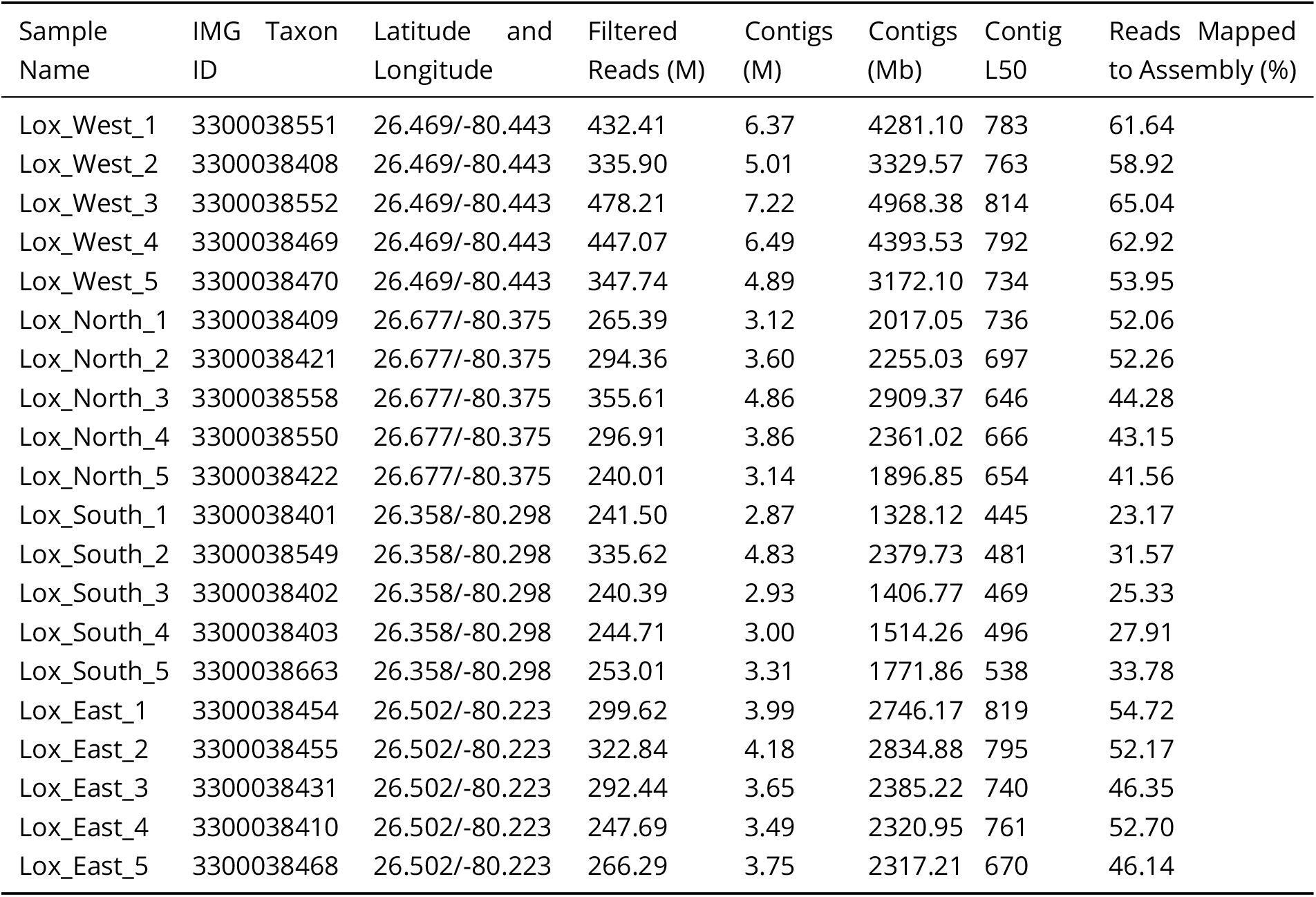
Sequencing and assembly statistics for 20 samples (4 sites, 5 replicates each) from the Loxahatchee Nature Preserve.

### Read prefiltering and assembly results

The target amount of raw sequence data was 45 Gb per sample (300M reads). The number of high quality raw reads per sample after quality trimming, filtering, artifact and contamination removal is shown in Table 1. While the replicates from Loxahatchee West were sequenced somewhat more deeply than other samples, there is no significant difference in the amount of sequence generated for the other three sites. The prefiltering and assembly modules of the workflow automatically generate several conventional measures of assembly quality that are provided in the README files and can be accessed via the JGI Data Portal. A subset of these measures, which helps with assessing the consistency of the samples and identifying the outliers and artifacts, is shown in Table 1. Despite the fact that the samples from Loxahatchee North, South, and East received a very similar amount of sequence, as shown in Figure 1a, assembly statistics indicate that the replicates collected at the South site differ from the rest. Box-and-whisker plots for the L50 metric (Fig. 1b, the smallest length of contigs for which the sum of lengths makes up half of the dataset size) and percent of reads mapped to the assembly (Fig. 1c) demonstrate that assemblies of South site replicates are significantly more fragmented, as indicated by much lower L50, and have fewer reads mapped to them. This may be due to the fact that the sediment at the South site has a large amount of sand, which hindered isolation of sufficient quantities of high-quality DNA (Jonathan B. Benskin, personal communication) thereby resulting in a suboptimal library and poor assembly. Variation of library quality due to the quality and quantity of the source DNA may not be immediately obvious with a functional and/or taxonomic analysis of unassembled reads but is prominently brought to the researcher’s attention by the DOE JGI Metagenome Workflow. It highlighted the differences between the South site and other sites due to the inconsistent performance of a sampling protocol, which may confound statistical analysis and obfuscate the true differences in functional and taxonomic composition.

**Figure 1.**
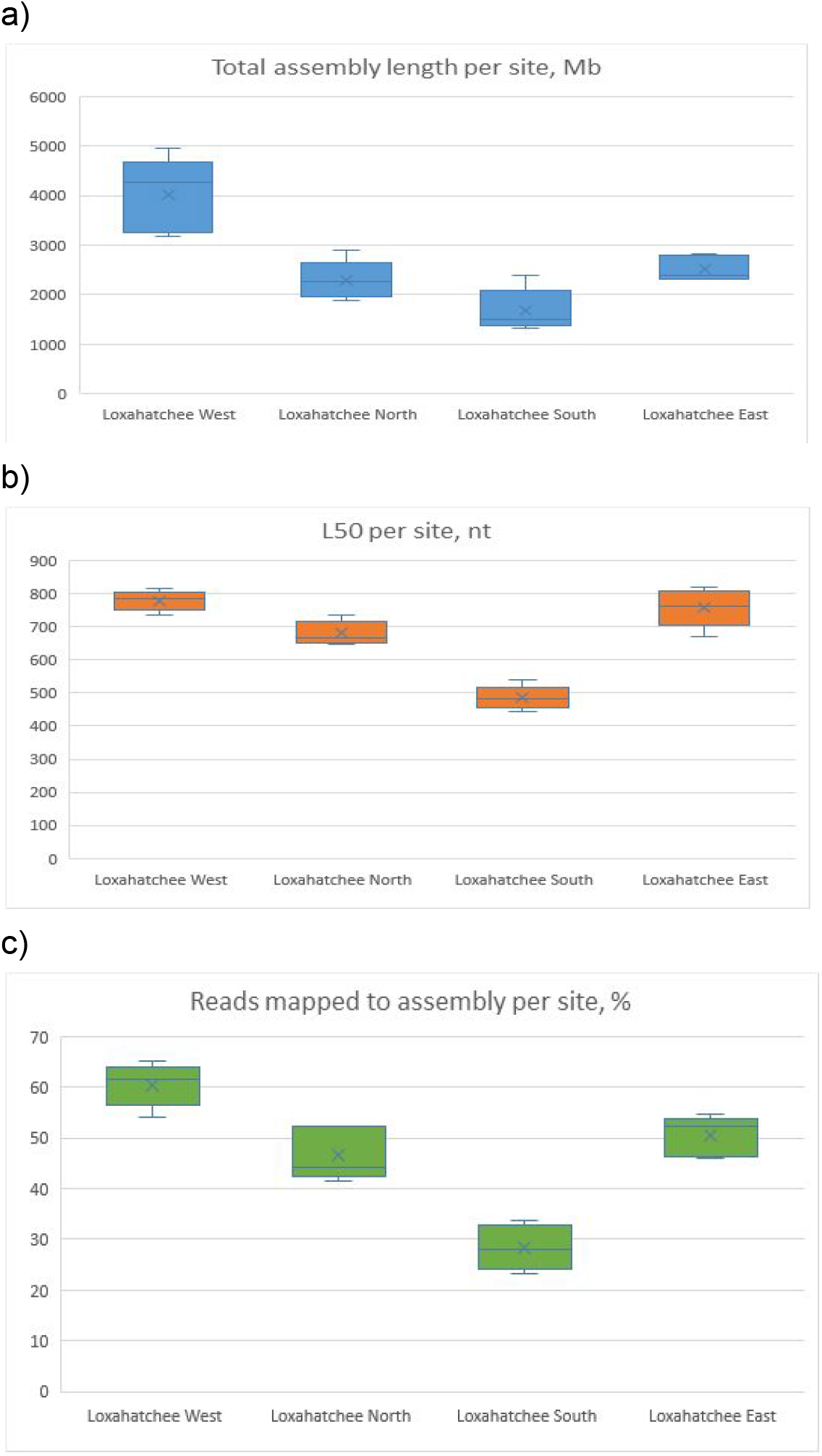
Box-and-whiskers plots of sequencing and assembly statistics for 4 sites in the Loxahatchee Nature Preserve. a) Total assembly length per site, Mb. b) L50 (the smallest length of contigs whose sum of lengths makes up half of the dataset size) per site, nt. c) Reads mapped to the assembly as percent of total number of reads generated for sample, per site, %.

### Annotation results

The DOE JGI Metagenome Workflow performs feature prediction (also known as structural annotation) on the assembled sequence and functional annotation of the predicted protein-coding genes (CDSs). Similar to the filtering and assembly modules, the annotation module generates summary statistics helpful for identification of artifacts and outlier samples. These statistics are provided in README files via the JGI Data Portal and can be also found in the IMG database on the Metagenome Details page of each dataset. A subset of the annotation measures for Loxahatchee samples is provided in Table 2. The results of functional annotation of CDSs appear to be highly consistent across the four sites, with 65.75+/-1.2% of all CDSs assigned to Clusters of Orthologous Genes (COGs)(11), 14.25+/-0.44% assigned to TIGRfams(12), 62.65+/-0.81% assigned to Pfams(13), and 40.2+/-1.85% assigned to KEGG Orthology (KO) Terms(14). However, the results of feature prediction summarized in Figure 2 paint a different picture. Again, the South site is different from the other three sites, having more predicted CDSs per Kb of assembled sequence (Fig. 2a), and a much higher number of predicted rRNAs per Mb of assembled sequence (Fig. 2b). Remarkably, there is no significant difference in tRNA counts (Fig. 2c). These observations are consistent with lower contiguity South site assemblies, as reflected in their lower L50 (Fig. 1b), which in turn results in fragmentation of longer protein coding genes, as well as long 16S/18S and 23S/28S rRNA genes. On the other hand, tRNAs, which are on average less than 100 nt long, are largely unaffected by the fragmentation of assembled sequences. Importantly, protein-coding genes, which span a large interval of sequence lengths, will be affected unevenly, with the copy number of longer proteins appearing to be higher in more fragmented assemblies, while shorter proteins will show no differences. These factors have to be taken into account when comparing functional composition of different samples and attempting to correlate it with various environmental factors. The feature prediction and functional annotation module of the DOE JGI Metagenome Workflow provides other indicators of the quality and consistency of metagenomic data: the counts of eukaryotic 18S and 28S rRNAs suggest the presence and abundance of eukaryotic genomes in the sample, which could derive from the eukaryotic members of the microbial community and/or host DNA in host-associated microbiomes. On the other hand, the relatively low percent of CDSs assigned to COGs and Pfams may indicate the presence of a large viral fraction in the community, since viral proteins are poorly represented in these protein and domain classification systems. All of these characteristics of the assembled metagenome need to be taken into account in comparative analyses, as they may affect the results of the taxonomic and functional annotation of the communities.

**Figure 2.**
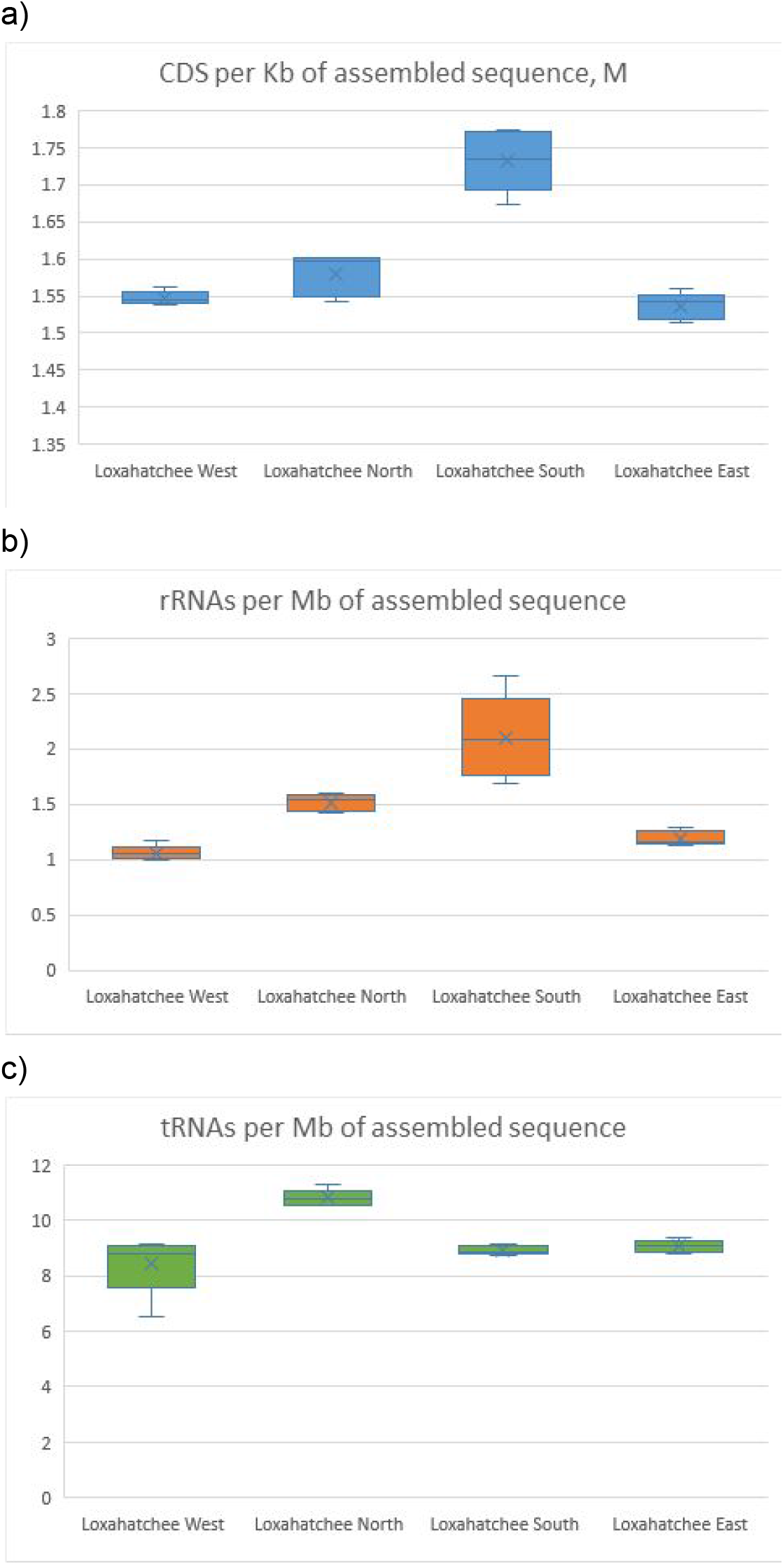
Box-and-whisker plots summarizing the results of structural annotation for 20 samples (4 sites, 5 replicates each) from the Loxahatchee Nature Preserve. a) Number of predicted CDSs per Kb of assembled sequence, millions. b) Number of predicted rRNA genes per Mb of assembled sequence. c) Number of predicted tRNA genes per Mb of assembled sequence.

**TABLE 2.**
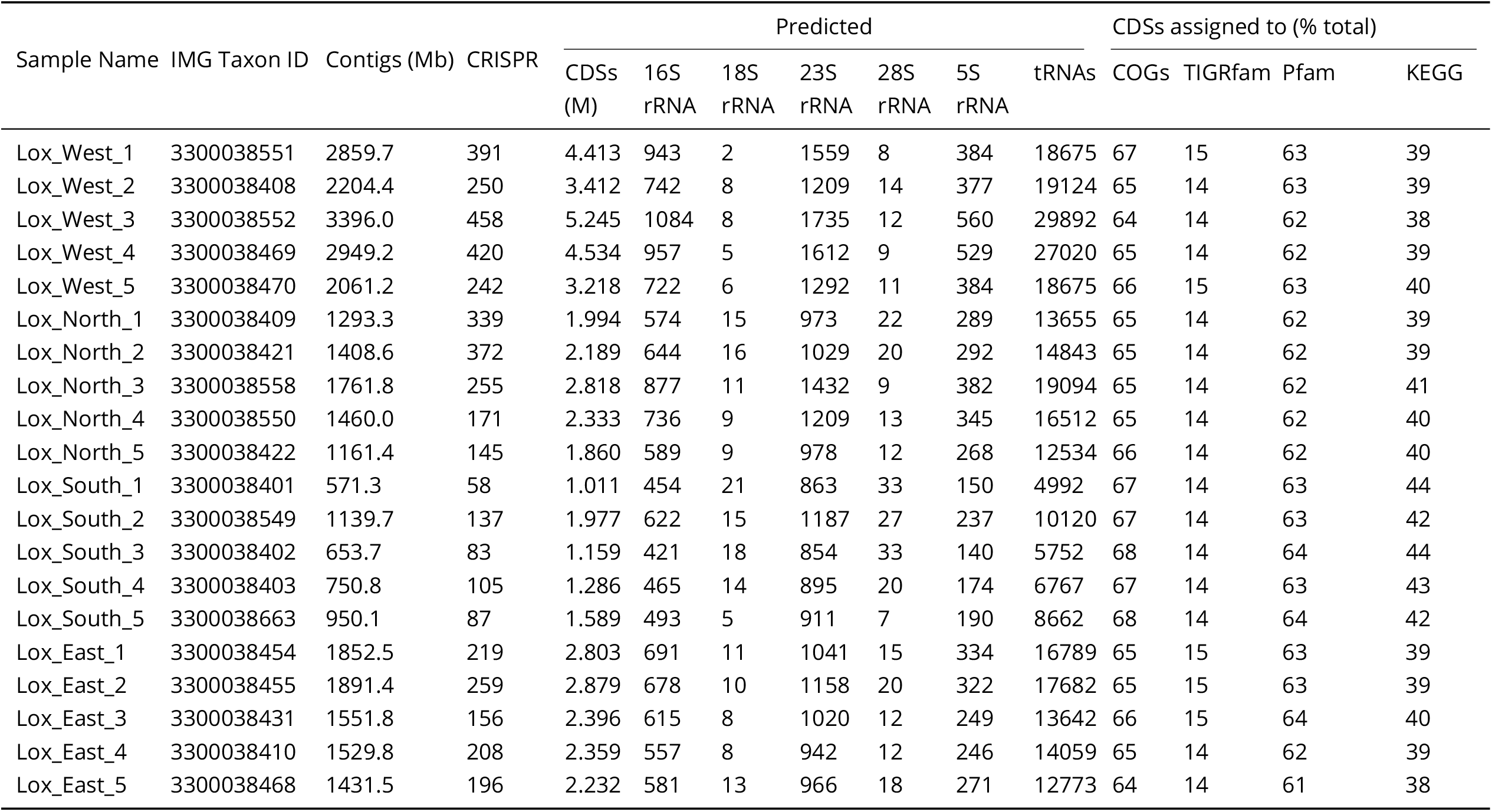
Annotation statistics for 20 samples (4 sites, 5 replicates each) from the Loxahatchee Nature Preserve.

### Binning results

The DOE JGI Metagenome Workflow includes automated binning of assembled sequences, as well as an initial characterization of bins in terms of completeness and contamination and quality. The bins are assigned to high-quality (HQ) and medium-quality (MQ) categories based on Minimum Information about a Metagenome-Assembled Genome (MIMAG) standards(15). Bins that do not meet the standards for HQ or MQ are discarded. For HQ and MQ bins additional data processing is performed: bins are assigned a predicted lineage based on the NCBI(16) and GTDB-tk(17) taxonomy. The results of genome binning for the Loxahatchee samples are summarized in Table 3. The vast majority of the bins generated for these datasets are MQ, and represent a minor portion of the total assembly typical of high-complexity metagenomes from soil and sediment samples. Binning results for each dataset can be accessed via the JGI Data Portal and in IMG, where a number of tools for search, analysis, and comparison of metagenome bins are available.

**TABLE 3.**
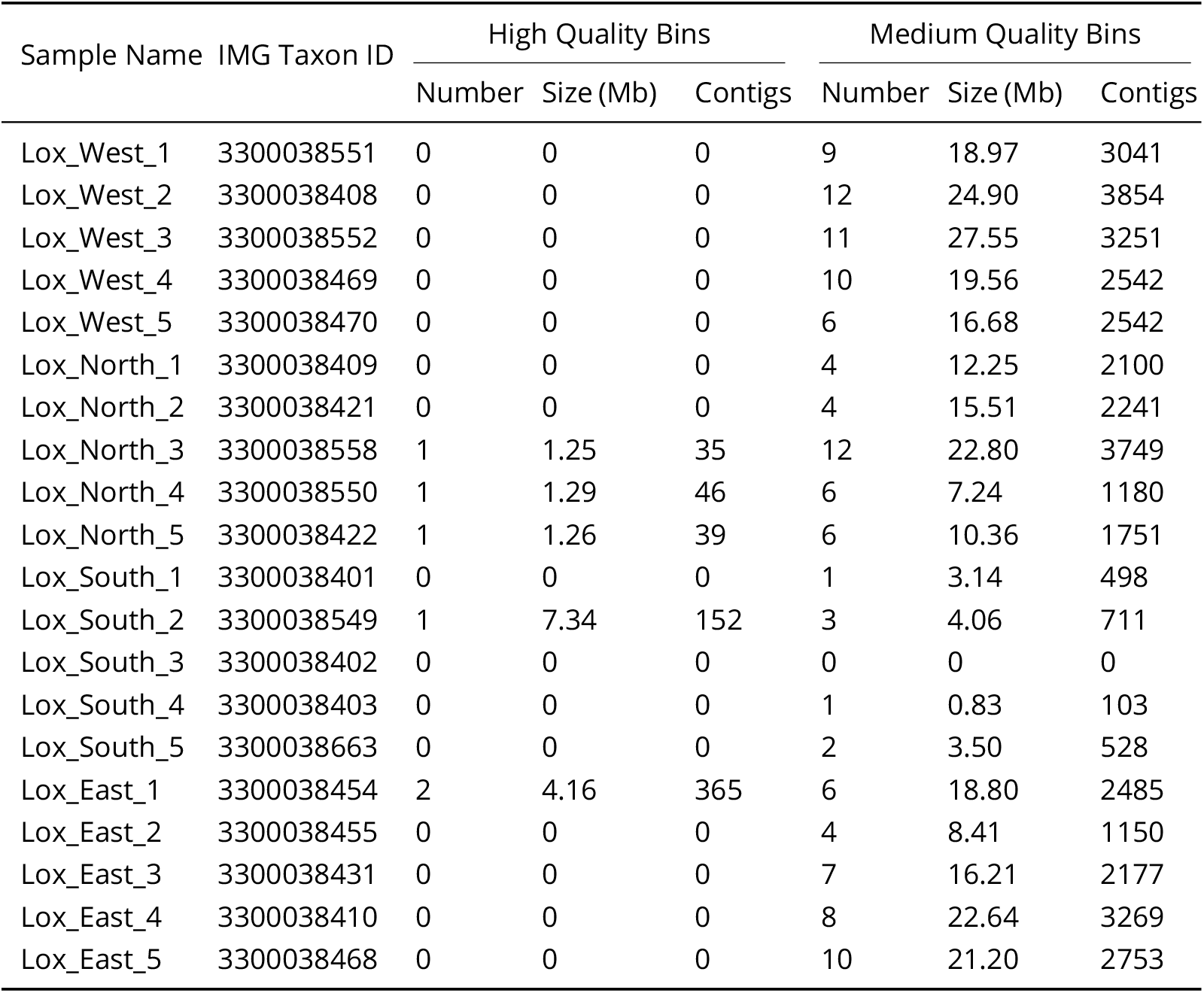
Binning statistics for 20 samples (4 sites, 5 replicates each) from the Loxahatchee Nature Preserve.

### Runtimes

We illustrate the typical computational requirements of the DOE JGI Metagenome Workflow on 20 samples from Loxahatchee Nature Preserve in Table 4. Filtering used Intel Xeon Gold 6140 processors using 32 vCPU and 324GB of RAM. For error correction, assembly, and mapping a mix of configurations was used. Some datasets were run on Intel Xeon Platinum 8000 series processors with different amounts of memory depending on the stage (16 vCPU and 128 GB of RAM for error correction, 64 vCPU and 512 GB of RAM for assembly, 32 vCPU and 256 GB of RAM for mapping). For others Intel Xeon Gold 6140 processors were used with 72 vCPU, 1.5 TB of RAM, and 5 TB of local disk. Runtime Assembly in Table 4 represents CPU hours for filtering, error correction, assembly, and mapping. For annotation, assembled metagenomic sequences were split into 10 MB shards. The splitting is performed by a wrapper script for optimal utilization of the JGI compute infrastructure and is not required to run the workflow. These 10 MB shards were then processed in parallel with each shard running on its own 2.3 GHz Haswell processor node with 128 GB of RAM. Binning was run on 2.3 GHz Haswell processor nodes with 128 GB of RAM.

**TABLE 4.**
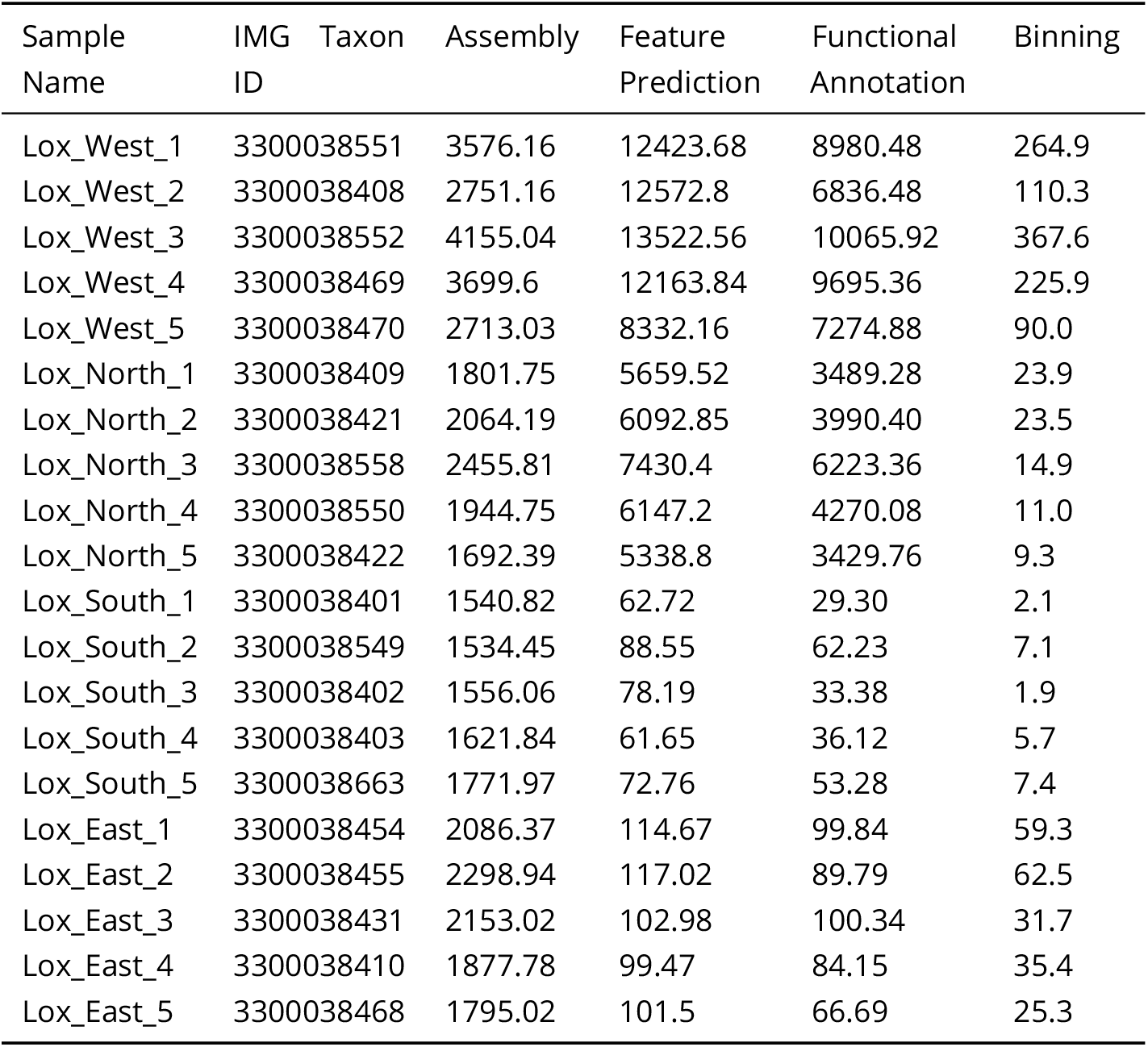
CPU hours for different modules in the JGI Metagenome Workflow on 20 samples from Loxahatchee Nature Preserve.

## DISCUSSION

The DOE JGI Metagenome Workflow provides automatic assembly, annotation and binning of metagenome datasets. It is largely based on publicly available software and databases supplemented with custom scripts and wrappers to control the workflow and to enable seamless integration of the input and output of different programs. Filtering, read correction, assembly, and mapping use a median of 2,004 CPU hours for current metagenomes such as the Loxahatchee sediment metagenomes, and can be performed on standard high-performance computing nodes such as Intel Xeon Platinum 8000 series processor with 256 GB memory. On average, the annotation module of the workflow (feature prediction, functional annotation and product name assignment) can process 1 million bp in 9 CPU hours on a 2.3 GHz Haswell processor (Intel Xeon Processor E5-2698 v3) node with 128 GB DDR4 2133 MHz memory. On the same Haswell node the entire binning workflow, from initial bin prediction, scaffold level cleanup, bin-level phylogenetic prediction, and estimation of contamination and completion, can process 100,000 scaffolds in an average of 13 CPU hours. The workflow modules for read filtering and metagenome assembly are available as a WDL file (https://code.jgi.doe.gov/BFoster/jgi_meta_wdl.git). The annotation and binning modules of the workflow are publicly available via the IMG system’s submission site (https://img.jgi.doe.gov/submit), which accepts assembled metagenome sequences in fasta format and requires submission of sample and project metadata as a condition of annotation and binning services. We plan to continue to improve the workflow by updating reference database versions, extending the existing software and adding new tools that allow the identification and characterization of more features in the metagenome datasets, as well as improving the performance by making changes geared towards exploiting the specific infrastructure the workflow is utilizing.

## MATERIALS AND METHODS

### Data input

Standard metagenomes at JGI currently use 100 ng of genomic DNA, sheared to 300 bp using the Covaris LE220 and size selected with SPRI using TotalPure NGS beads (Omega Bio-tek). The fragments are treated with end-repair, A-tailing, and ligation of Illumina compatible adapters (IDT, Inc) using the KAPA-HyperPrep kit (KAPA biosystems) to create an unamplified Illumina library which is then sequenced 2×150 bp on the Illumina NovaSeq 6000 using S4 flowcells. The workflow can be used on paired-end Illumina datasets; kmer sizes for assembly should be adjusted if reads are shorter than 150 bp.

### Sequence data preprocessing

Data is processed using Real-time Analysis (RTA) version3.4.4(https://support.illumina.com/downloads.html). BBDukversion38.79from the BBTools package (https://jgi.doe.gov/data-and-tools/bbtools/) is used to remove contamination, trim reads that contain adapter sequence and quality trim reads where quality drops to 0. Furthermore, it is used to remove reads that contain 4 or more ’N’ bases, have an average quality score across the read of less than 3 or have a minimum length less than or equal to 51 bp or 33% of the full read length. Homopolymer streches of 5 Gs or more at the ends of reads are removed. Reads that can be mapped with BBMap from BBTools to masked human, cat, dog and mouse references at 93% identity are separated into a “chaff” file and not used in assembly. In an abundance of caution, reads aligned to common microbial contaminants described in the literature such as *Ralstonia pickettii* and *Acinetobacter calcoaceticus*(18, 19, 20, 21) are also separated into a “chaff” file. Masked references can be found at https://portal.nersc.gov/dna/microbial/assembly/bushnell/fusedERPBBmasked2.fa.gz. For convenience chaff files are provided on JGI’s data portal.

### Assembly

Filtered reads are error corrected using bbcms version 38.44 from BBTools with a minimum count of 2 and a high count fraction of 0.6. Bbcms uses a count-min sketch to store kmer counts, making it a scaleable solution for error correction of metagenomic datasets. For computational efficiency, interleaved fastq files are split into two separate files. These split error corrected files are assembled with metaSPAdes version 3.13.0 using the “metagenome” flag, running the assembly module only (i. e. without error correction) with kmer sizes 33,55,77,99,127. Contigs that are smaller than 200 bp are discarded. Filtered reads are mapped back to contigs larger than 200 bp using BBMap 38.44 with “interleaved” as true, “ambiguous” as random, and “covstats” option specifying a contig coverage file for subsequent analysis of abundance of various populations and genes. The coverage file contains information on average fold coverage, length, GC content, percent of bases covered, number of reads by strand, read GC, median fold and standard deviation of coverage.

### Feature prediction

The assembled contigs are passed on to the annotation module of the workflow, which first predicts non-coding RNA genes (tRNAs, rRNAs and other RNAs), followed by the identification of Clustered Regularly Interspaced Short Palindromic Repeats (CRISPR) and protein coding genes (CDSs) as shown in Fig. 3a. Prediction of tRNAs is performed using tRNAscan-SE 2.0.6(22) in “bacterial” and “archaeal” search modes. This allows the workflow to select the best annotation mode and ensure higher annotation accuracy for metagenomic contigs of different taxonomic origin, since many archaeal tRNAs cannot be predicted in “bacterial” or “general” modes. For each contig the number of tRNAs with known isotype returned by each mode is compared. The results from the mode with the higher number of tRNAs with known isotype get reported and if both modes have returned the same number, the results from the “bacterial” mode are included in the final annotation. Ribosomal RNA genes (5S, 16S, 23S) as well as other non-coding RNA genes (ncRNAs) including tmRNA, antisense RNAs, etc. and RNA regulatory features, such as various binding sites and motifs (“misc_bind”, “misc_feature”, “regulatory”) are identified by comparing the contigs via cmsearch from the INFERNAL 1.1.3 package(23) against the Rfam 13.0 database(24) using the trusted cutoffs parameter (-cut_tc). If any reported hits are overlapping even by 1 bp and they belong to the same Rfam class, the lower scoring of the two is discarded. CRISPR elements are identified using a version of CRT-CLI 1.2 modified in-house as described previously(25). For the search parameter the minimum and maximum repeat lengths are set to 20 and 50 bp, respectively, whereas the minimum and maximum spacer length is set to 20 and 60 bp, respectively. The search window size is set to 7 bp and an element needs to have at least three repeats to get reported. Protein-coding genes are predicted via a combination of Prodigal 2.6.3(26) and GeneMarkS-2 1.07(27). Prodigal is executed in “meta” mode and with the ‘-m’ argument so that genes won’t be built across runs of Ns. GeneMark is run with ‘-Meta mgm_11.mod’ and ‘-incomplete_at_gaps 30’. CDS shorter than 75 bp (25 amino acids) are discarded. The last step of the feature prediction combines the results from all tools and attempts to resolve overlaps between features of different types. Two features are considered to overlap if they share more than 10 bp or more than 90 bp in the case of two CDSs. The regulatory RNA features (misc_bind, misc_feature and regulatory) are allowed to overlap with any other feature type. In case of an overlap between other types of features the lower-ranked feature gets removed. The feature ranking order is rRNA > tRNA > ncRNA, tmRNA > CRISPR > GeneMarkS-2 > Prodigal. Before deleting a CDS that overlaps with another feature over its 5’ end, first an attempt is made to find an alternative start site for the protein-coding gene that removes the overlap. Functional annotations of RNA features are based on their descriptions provided by the tool or database used to predict them: tRNA isotype (amino acid and codon) as well as potential pseudogene annotation is provided by tRNAscan-SE, while product names for rRNAs, ncRNAs and regulatory RNA features are derived from the corresponding Rfam models. Functional annotation and product name assignment for protein sequences of the non-overlapping CDSs is performed by the functional annotation module.

**Figure 3.**
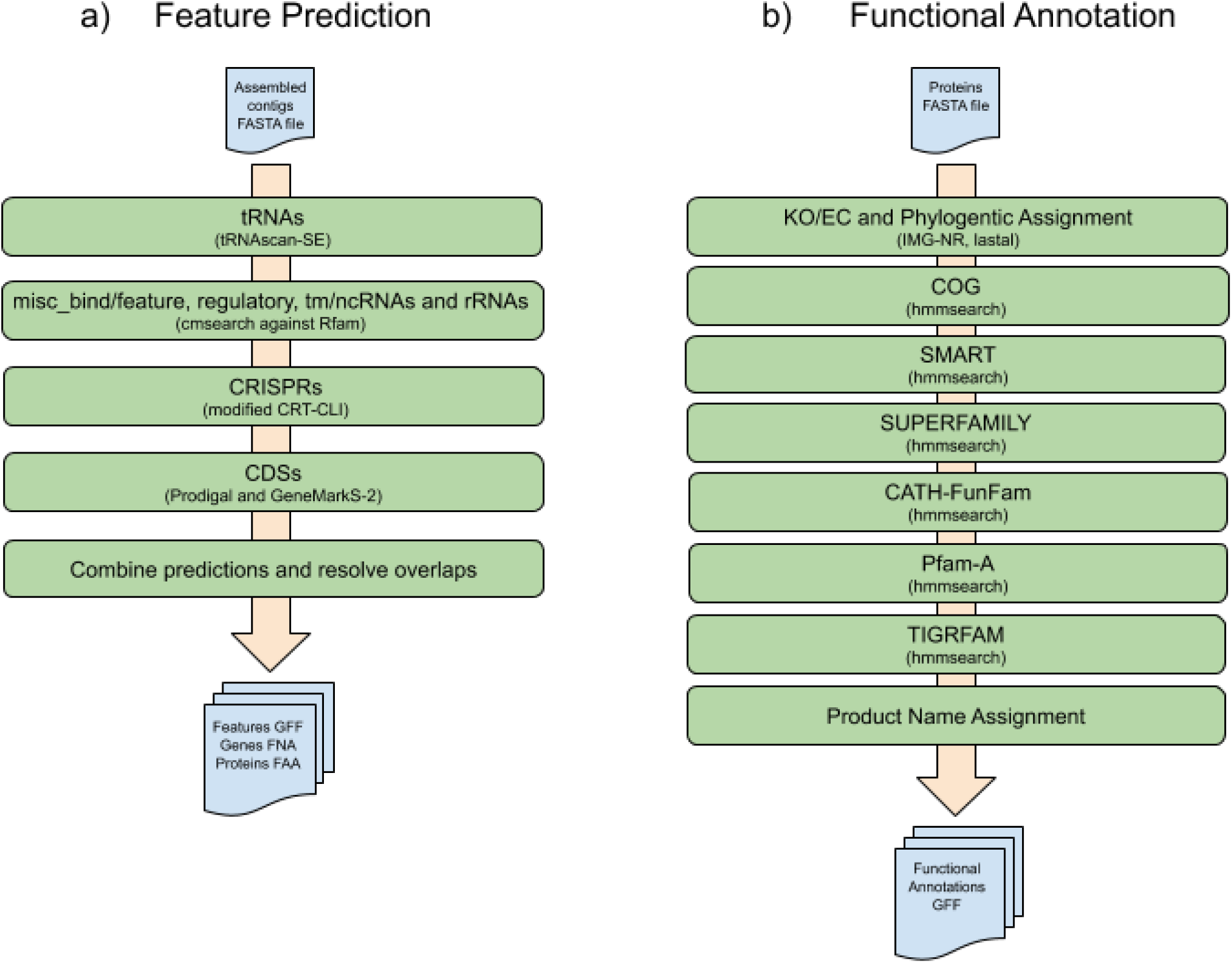
Workflow of a) Feature Prediction and b) Functional Annotation

### Functional annotation

Functional annotation for metagenomes consists of associating protein-coding genes with KO terms, Enzyme Commission (EC) numbers, COG assignments, SMART domains, SUPERFAMILY assignments, CATH-FunFam annotations, Pfam and TIGRFAM annotations as shown in Fig. 3b. Genes are associated with KO terms and EC numbers based on results of sequence similarity search of metagenome proteins against a reference database of isolate proteomes using lastal 1066 from the LAST package(28) with default parameters. The reference database of isolate proteomes (IMG-NR) is composed of all non-redundant protein sequences encoded by public, high quality genomes in the current version of the IMG database. For each metagenome protein the top five LAST hits are considered. At least two of the top five hits need to have a KO assignment and all hits that have a KO assignment need to list the same combination of KO terms. If both conditions are met, the same combination of KO terms is assigned to the query gene if the alignment length for any of the hits with KO assignment covers at least 70% of the shorter one of query and subject. Proteins are associated with COGs by comparing protein sequences to the COG Hidden Markov Models (HMMs) created from the updated 2014 models using HMMER 3.1b2(29), and a thread-optimized version of hmmsearch(30), with a per-domain e-value cutoff (-domE) of 0.01. Since an alignment of a protein to the model may be fragmented, i. e. there may be multiple aligned segments of the two, these are concatenated and their cumulative alignment length calculated. If the cumulative align-ment length is less than 70% of the shorter of the two (the protein or the model), such a hit is discarded. In addition, if a protein has hits to different COG models and their alignments overlap significantly (by more than 10% of the length of the shorter model), the hit to the model with the lower full sequence bitscore is discarded; for significantly overlapping hits with the same bit score, the hit with the higher e-value gets removed. The same thread-optimized version of hmmsearch, as well as parameters, filtering and overlap resolution rules are used to assign protein sequences to the 01_06_2016 version of the SMART database(31), the 1.75 version of the SUPERFAMILY database(32) and the frozen set of the 4.2.0 version of the CATH-FunFam database(33). Proteins are associated with Pfam-A by comparing protein sequences to version 30 of the Pfam database using thread-optimized version of hmmsearch from HMMER 3.1b2. Model-specific trusted cut-offs are used with (-cut_tc option in hmmsearch) and for overlapping hits that belong to the same Pfam clan the lower scoring one is removed. Proteins are associated with TIGRFAMs using version 15.0 of the TIGRFAM database and hmm-search with a per-domain e-value cutoff (-domE) of 0.01. All hits that don’t cover at least 70% of the shorter of the protein or model get discarded. Furthermore if two hits overlap for more than 10% of the length of the shorter model, the hit to the lower scoring model (by bit score) is discarded. Protein product names are assigned based on the name of their associated protein families in the order of priority KO term > TIGRFAM > COG > Pfam. If multiple TIGRFAMs with different isology types are associated with a protein, only one TIGRFAM is assigned in the order equivalog > hypoth_equivalog > paralog > exception > equivalog_domain > hypoth_equivalog_domain > paralog_domai n> subfamily > superfamily > subfamily_domain > domain > signature > repeat. Proteins without any of the above assignments are annotated as “hypothetical protein”. Proteins associated with multiple protein families of the same type (KO term, TIGRFAM, COG or Pfam) are annotated with a product name consisting of concatenation of individual protein family names joined with “/”. Multiple repetitions of the same protein family are collapsed to a single instance. The contig coverage information is used to calculate so-called “estimated gene copies”, whereby the number of genes in a certain group, such as a COG or Pfam protein family, is multiplied by the average coverage of the contigs from which these genes were predicted. This step is important for accurate estimation of abundance of protein families and takes into account different abundance of populations found in the assembled metagenome sequences.

### Taxonomic annotation

For taxonomic annotation of metagenomes the best LAST (28) hits of CDSs computed as described above for KO term assignment are used. The taxonomy of the best hit is assigned to each metagenome protein. The taxonomy of metagenome contigs (“scaffold lineage”) is predicted based on the majority rule, whereby the lineage at the lowest taxonomic rank to which at least 50% CDSs encoded by the metagenomic contig have hits is assigned. Similar to protein family annotations, contig coverage information is used to estimate the abundance of various lineages in the community by multiplying contig counts by their average coverage.

### Binning

The assembled contigs and coverage file generated per metagenome is used as input to the MetaBAT v2.12.1(34) program to generate genome bins based on the consistency of coverage and tetranucleotide frequency. The genome bins then undergo contamination removal, wherein the per scaffold phylum information generated by the annotation module (“scaffold lineage”) is used to remove scaffolds per bin that are not assigned to the predominant phylum. The post-processed bins are fed to the CheckM v1.0.12(35) program to determine genome completion and contamination estimates. These estimates along with the per scaffold rRNA and tRNA information generated by the annotation module, is used to assign HQ or MQ value to each bin, per MIMAG standards. The HQ and MQ bins are then subject to phylogenetic lineage determination by two methods. First, an internal IMG program computes the phylogenetic lineage per genome bin using the per scaffold lineage generated by the annotation module. Next, the GTDB-tk v0.2.2 program computes per bin lineage by placing them into domain-specific, concatenated protein reference trees. The high and medium quality bins, along with the corresponding data processing metadata, are loaded into IMG for user access and download.

### Pre-formatted tables

To assist with preparing publications 9 tables are generated. Information on what is contained in each table is described in Table 5.

**TABLE 5.** Preformatted tables

| Table number | Table information |
| --- | --- |
| 1 | Study information |
| 2 | Sample information |
| 3 | Library information |
| 4 | Sequence process |
| 5 | Assembly statistics |
| 6 | Annotation parameters |
| 7 | Functional diversity |
| 8 | Metagenome properties |
| 9 | Taxonomic composition |

## Availability of data and materials

The metadata for these samples can be found in GOLD (https://gold.jgi.doe.gov/) using GOLD study Gs0136122. Raw reads, as well as intermediate results and final assembly and annotation data can be found in the JGI Data Portal (https://genome.jgi.doe.gov) by following links from the GOLD study or by using IMG taxon identifiers provided in Table 1. A WDL for filtering and genome assembly (v1.0) is available at https://code.jgi.doe.gov/BFoster/jgi_meta_wdl.git. IMG for annotation (v5.0.19) and binning (v1.0) is available at https://img.jgi.doe.gov/.

## ACKNOWLEDGMENTS

The work conducted by the U.S. Department of Energy Joint Genome Institute, a DOE Office of Science User Facility, is supported by the Office of Science of the U.S. Department of Energy under Contract No. DE-AC02-05CH11231. We thank the Advanced International Certificate of Education (AICE) Biology class at Florida’s Boca Raton Community High School and Mr. Jonathan Benskin for partnering with the JGI on the Everglades metagenome studies.

## Notes

https://gold.jgi.doe.gov/

https://code.jgi.doe.gov/BFoster/jgi\_meta\_wdl.git

https://img.jgi.doe.gov/

### Competing Interest Statement

The authors have declared no competing interest.

